# Co-folding of Membrane Proteins and Lipid Molecules Improves Membrane-Protein Structure Prediction Accuracy

**DOI:** 10.64898/2026.03.24.713847

**Authors:** Hiroaki Oheda, Masao Inoue, Toru Ekimoto, Tsutomu Yamane, Mitsunori Ikeguchi

## Abstract

Recent advances in deep-learning-based structure prediction have greatly improved the accuracy of protein structure modeling and enabled prediction of biomolecular complexes. However, the surrounding molecular environments are typically not represented explicitly, and their effects are therefore captured only implicitly. This limitation is particularly relevant for membrane proteins, whose structures and interactions are strongly influenced by the membrane environment. Here we introduce co-folding of membrane proteins and lipid molecules (CoMPLip), a method that imposes an explicit membrane context by co-folding proteins with lipid molecules during AlphaFold 3 (AF3) predictions. In CoMPLip, lipid molecules arrange into bilayer-like configurations around transmembrane regions, providing a membrane-like environment during structure prediction. Across benchmark datasets of ligand-bound membrane proteins, full-length single-pass membrane proteins, and dynamic transporters, CoMPLip improves ligand-pose prediction, promotes correct extracellular-intracellular domain separation, and enables sampling of multiple conformational states. CoMPLip is training-free and is compatible with the existing AF3 workflows.

## Introduction

Membrane proteins mediate a wide range of essential biological processes, including signal transduction, transport, and energy conversion, and constitute a large fraction of drug targets ^1^. Determining their three-dimensional structures is therefore crucial for understanding molecular mechanisms at atomic resolution and enabling structure-based drug design (SBDD).

Recent advances in deep learning–based structure prediction have transformed structural biology. Methods such as AlphaFold2 (AF2) ^2^ have enabled the highly accurate protein structure prediction directly from sequence, including many membrane proteins ^3–5^. More recently, next-generation models such as RoseTTAFold All-Atom ^6^, AlphaFold 3 (AF3) ^7^, Boltz-2 ^8^, and Chai-1 ^9^ have extended structure prediction to biomolecular complexes involving nucleic acids, small molecules, and other macromolecules, further broadening the applicability of AI-based structural modeling. Despite these advances, accurate prediction of membrane-protein structures remains challenging in several situations. For example, in full-length single-pass membrane proteins, individual domains are often predicted accurately. However, extracellular and intracellular domains can be incorrectly placed in direct contact across the transmembrane region ^10^. In ligand-bound membrane proteins, the predicted ligand poses can deviate substantially from the experimentally observed binding modes ^11^. Additionally, for transporters and other dynamic membrane proteins, structural predictions often capture only a single conformation, even when multiple functional states are known experimentally ^12,13^.

A possible reason for these limitations is that the membrane environment is not explicitly represented in current prediction protocols. Although training datasets contain many membrane-protein structures determined in membrane-mimetic environments, lipid bilayers themselves are not explicitly modeled during prediction. Because membrane proteins function within lipid bilayers, the absence of an explicit membrane context may reduce the physical realism of predicted structures and limit prediction accuracy in certain cases ^14^.

Here we introduce co-folding of membrane proteins and lipid molecules (CoMPLip), a simple strategy for incorporating an explicit membrane environment into AlphaFold-based structure prediction. In CoMPLip, lipid molecules are provided as additional molecular inputs during AF3 prediction, allowing the protein and surrounding lipids to be predicted simultaneously. The lipid molecules spontaneously organize around the transmembrane region, forming bilayer-like arrangements that impose a membrane-like structural context during prediction.

We demonstrate that CoMPLip improves prediction performance in three representative challenges of membrane protein structure prediction: (i) accurate placement of ligands in ligand-bound membrane proteins, (ii) correct spatial separation of extracellular and intracellular domains in full-length single-pass membrane proteins, and (iii) sampling of multiple conformational states in dynamic membrane proteins. We further analyze the influence of lipid parameters, including lipid copy number and chain length, and introduce a CoMPLip-specific ranking score (*S*_CoMPLip_) that focuses on the prediction quality of the target protein–ligand complex, while excluding contributions from added lipids.

## Results and Discussion

### Improved ligand-binding pose prediction with CoMPLip

First, we addressed challenge (i) defined in the Introduction: accurate prediction of ligand-binding poses in membrane proteins. To evaluate whether CoMPLip improves ligand-pose accuracy, we examined the complex of the *Escherichia coli* regulator of sigma-E protease (RseP) bound to the inhibitor batimastat (BAT) as a representative example, and compared the predicted structures with the experimental structure (PDB ID: 7W6X) ^15^.

Predictions were performed under two conditions: standard AlphaFold 3 (AF3) prediction without lipids, and CoMPLip prediction with lipids. Under CoMPLip conditions, 100 molecules of 1-monoolein, the crystallization lipid used for the 7W6X structure, were included as co-folded molecules (Fig. 1). For each condition, predictions were carried out using five seeds, generating 25 models, and the representative model was selected according to the standard AF3 ranking score. To quantify prediction accuracy, the predicted structures were aligned to the experimental structure using Cα atoms in the transmembrane (TM) region (residues 2–33, 94–122, 376–415 and 423–447 defined in Ref. 15), and root-mean-square deviations (RMSDs) were calculated for both the TM region and BAT. The TM-region RMSD indicated similarly accurate protein structures in both conditions (0.58 Å without lipids and 0.74 Å with lipids). In contrast, the ligand RMSD differed substantially, decreasing from 10.79 Å without lipids to 1.37 Å with lipids. These results indicate that CoMPLip markedly improves the accuracy of the predicted RseP–BAT binding pose.

**Figure 1.**
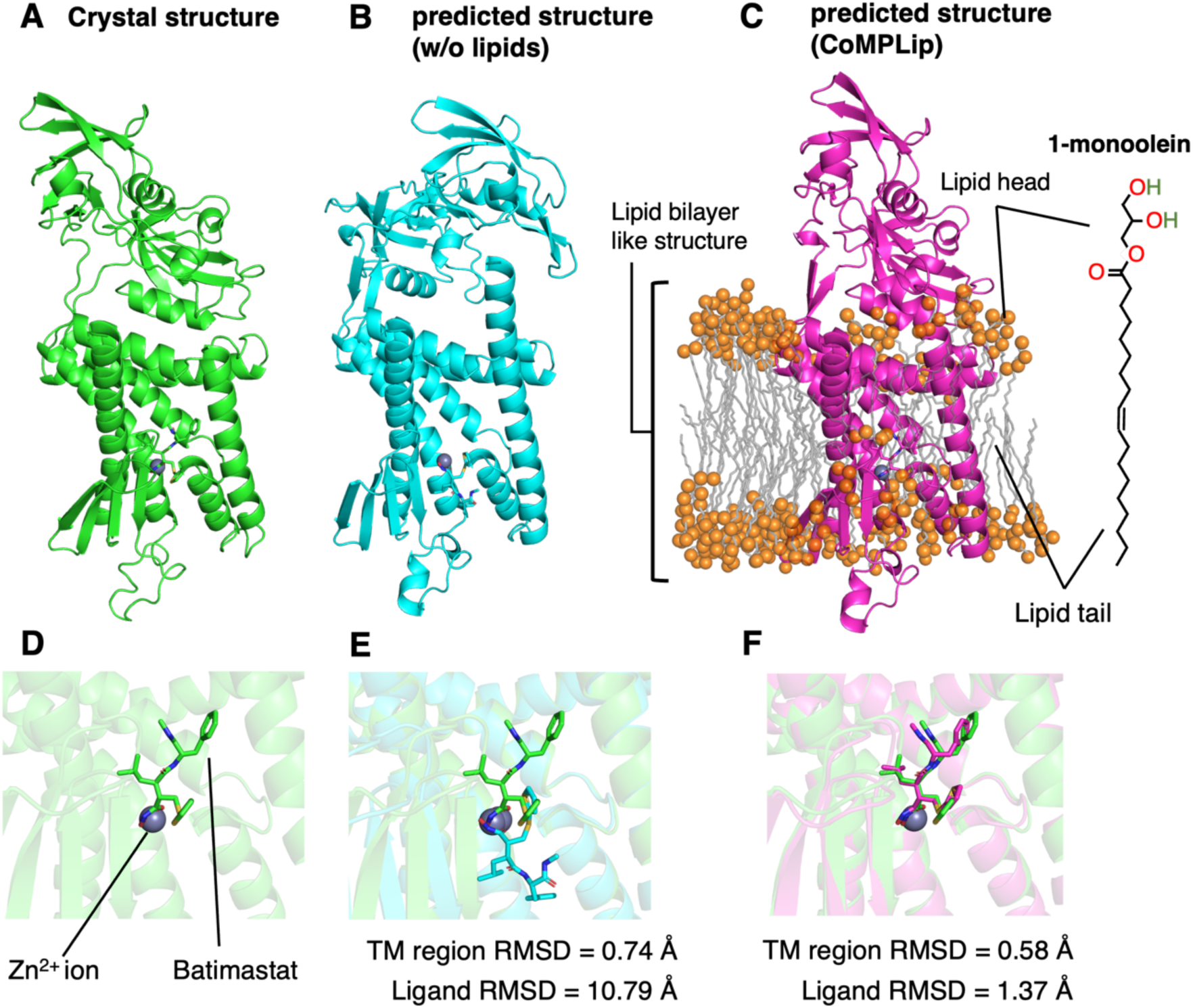
Experimental and predicted structures of the RseP–BAT complex. **A** Experimental structure of RseP with Batimastat (BAT) and Zn²⁺ (PDB ID: 7W6X; green). **B** AF3 prediction of the RseP–BAT complex without (w/o) lipids (cyan). The model with the highest AF3 ranking is also shown. **C** CoMPLip prediction with 100 molecules of 1-monoolein (magenta). The model with the highest AF3 ranking is also shown. Predicted structures were aligned to the experimental structure using Cα atoms in the transmembrane (TM) region, and the RMSDs were calculated for the TM region and BAT. **D–F** The RMSDs were calculated as described above.

In the CoMPLip prediction, the added lipid molecules spontaneously organized into a bilayer-like arrangement surrounding the transmembrane region of RseP (Fig. 1C), providing a membrane-like environment during structure prediction.

To further investigate the effect of CoMPLip on ligand-pose prediction accuracy in the RseP–BAT complex, the number of prediction seeds for each prediction condition was increased to 100, yielding 500 models, and statistical analysis was performed using the resulting predicted structures. Ligand-binding poses were classified as correct or incorrect according to the RMSD of the ligand relative to the experimental structure (PDB ID: 7W6X). Poses with a ligand RMSD < 4 Å were classified as correct. Under this classification, 114 of 500 models were correct without lipids, whereas 254 of 500 models were correct with lipids (Fig. 2). To further assess the structural effects of CoMPLip on RseP, we measured the RMSD of the MREβ region, an intramembrane region important for RseP activity ^16^, relative to the experimental structure. The average MREβ RMSD across the predicted structures decreased from 2.97 Å without lipids to 1.04 Å with lipids. In the CoMPLip predictions, multiple lipid molecules were consistently positioned near hydrophobic residues in the MREβ region (Supplementary Fig. 1). Across all predicted structures, models generated with CoMPLip were concentrated in the region of low MREβ RMSD values (Supplementary Fig. 2). Together, these results suggest that CoMPLip improves the structural accuracy of the MREβ region, which likely contributes to more accurate ligand-pose prediction.

**Figure 2.**
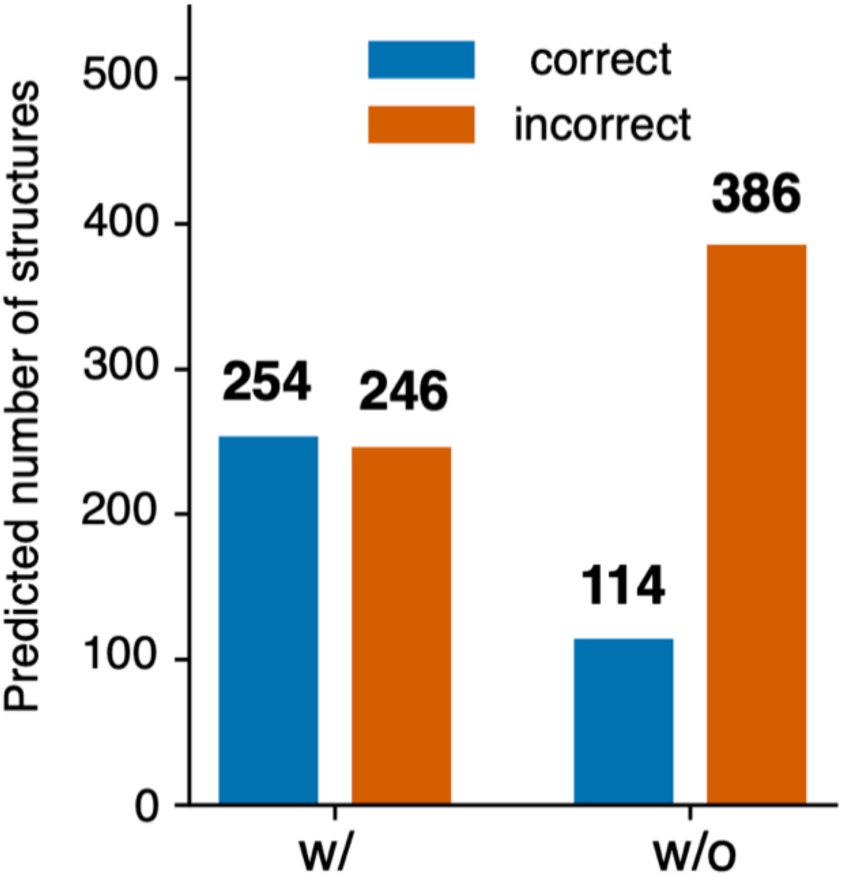
Ligand-binding pose prediction accuracy. Among the 500 predicted models per condition (standard AF3 without lipids or CoMPLip with lipids), ligand poses were classified as correct when the BAT RMSD relative to the experimental pose (PDB ID: 7W6X) was < 4 Å. Correct poses were obtained for 114/500 models without lipids and 254/500 models with 100 molecules of 1-monoolein.

### Optimization of CoMPLip parameters

In addition to the input parameters for the standard AF3 prediction, CoMPLip requires parameters that specify the added lipid molecules. In particular, lipid carbon chain length and lipid copy number are expected to influence prediction accuracy. To examine their effects, the RseP–BAT system was used as a test case. Prediction accuracy was evaluated using the ligand RMSD of BAT relative to the experimental pose (PDB ID: 7W6X ^15^). Predictions were considered successful when the ligand RMSD was < 4 Å.

The effect of lipid carbon chain length was examined using a series of 1-monoacylglycerols (general structure: R–C(=O)O–CH₂–CH(OH)–CH₂OH; *n* = 1–17, where *n* denotes the number of carbons in the alkyl chain R and R = CH₃(CH₂)*_n_*_−1_). For each value of *n*, CoMPLip predictions were performed using 100 lipids, and the representative structure was selected using the conventional AF3 ranking score. A lipid-free condition is shown in Fig. 3A. Ligand RMSD values < 4 Å were consistently obtained for *n* ≥ 14 (Fig. 3A). The effect of lipid copy number was also examined by varying the number of 1-monoolein molecules from 0 to 100 in steps of 10. Ligand RMSD values < 4 Å were obtained when more than 80 lipid molecules were included (Fig. 3B). In the RseP system, prediction accuracy improved with increasing lipid copy number and with longer lipid carbon chains. These results provide practical guidance for parameter selection. 1-monoolein (glycerol monooleate) contains an oleoyl chain (C18:1). Since the predominant lipid carbon chain length in the E. coli inner membrane, where RseP is localized, are C16–C18 ^17^, the C18 chain length of 1-monoolein represents a reasonable starting point when selecting lipids for CoMPLip. In addition, a sufficient number of lipid molecules is required to adequately cover the hydrophobic transmembrane surface. Approximately 80 lipid molecules provide practical guidelines for the four-pass transmembrane protein, RseP. Proteins with larger transmembrane surfaces may require proportionally more lipid molecules, whereas single-pass transmembrane proteins may require fewer molecules. This was further examined in a later section using a single-pass transmembrane protein.

**Figure 3.**
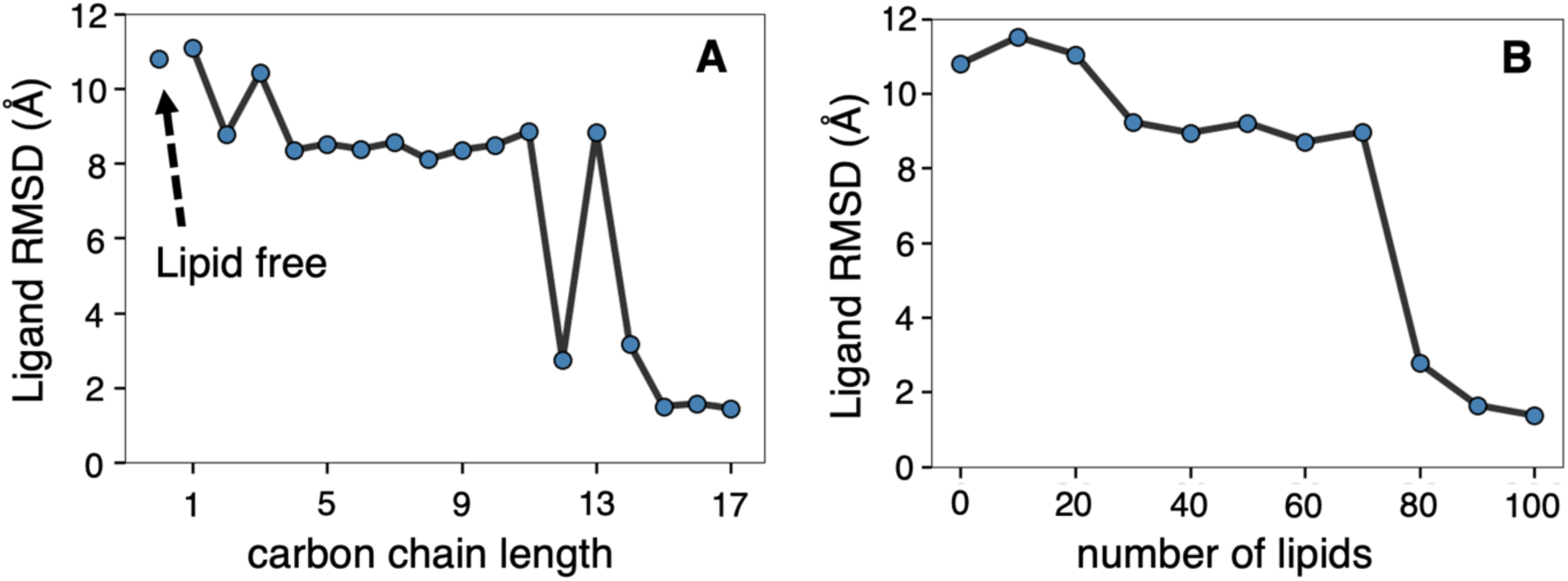
Effects of lipid number and carbon chain length on CoMPLip predictions of the RseP–BAT complex. Ligand (BAT) RMSD values were calculated after aligning predicted structures to the experimental reference (PDB ID: 7W6X) using Cα atoms of transmembrane (TM) helices. **A** Effect of lipid carbon chain length. CoMPLip predictions were performed using a series of 1-monoacylglycerols (general structure: R–C(=O)O–CH_2_–CH(OH)–CH_2_OH; *n* = 1–17, where *n* denotes the number of carbons in the alkyl chain R and R = CH_3_(CH_2_)*_n_*_−1_). Lipid-free conditions are shown separately. **B** Effect of lipid copy number. The CoMPLip predictions were performed with 1-monoolein copy numbers ranging from 0 to 100 in steps of 10.

### Lipid-independent protein structure ranking score

When CoMPLip was applied, the standard AF3 ranking score was influenced by the large number of added lipid molecules, which often received low confidence scores. As a result, the standard score may underestimate the accuracy of the target protein–ligand structure. To address this issue, we defined a new ranking score for CoMPLip predictions that focuses on the target protein and ligand while excluding contributions from the added lipids. The standard AF3 ranking score (S_AF3_) is defined as ^7^

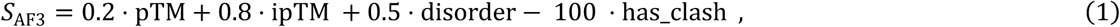

where “pTM” is the predicted TM-score, “ipTM” is the predicted interfacial TM-score, “disorder” is a scalar in the range of 0–1 indicating the fraction of the prediction structure that is disordered, and “has_clash” indicates whether the predicted structure contains severe atomic clashes. In CoMPLip predictions, the total pTM value reflects contributions not only from the protein and ligand but also from the added lipid molecules, and the ipTM value includes interfaces involving protein–ligand, protein–lipid, ligand–lipid, and lipid–lipid interactions. Because lipid-related terms are more numerous than those associated with the target protein and ligand, they can obscure the structural accuracy of the protein–ligand complex. To mitigate this effect, we defined a new ranking score for CoMPLip predictions (*S*_CoMPLip_) using only the protein and ligand components of the AF3 output, as follows:

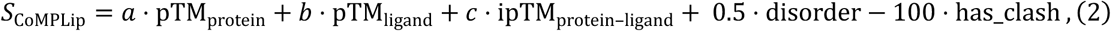

where “pTM_protein_” and “pTM_ligand_” are the pTM scores of the protein and ligand, respectively, and “ipTM_protein–ligand_” represents the interfacial score between the protein and ligand. Parameters *a*, *b*, and *c* are non-negative weights. Here, “ligand” refers to the target small-molecule ligand of interest and excludes the lipid molecules added in CoMPLip (for example, BAT in the prediction of the RseP–BAT complex).

Parameters *a*, *b,* and *c* were optimized using a grid search (step size 0.1) to maximize the correlation with the ligand RMSD across the 500 structures predicted for RseP–BAT complex (the dataset used in Figs. 2 and 3). The optimal weights are *a* = 0.2, *b* = 0.7, *c* = 0.1, yielding:

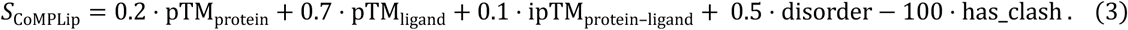

Because these coefficients were optimized for a single system (RseP–BAT), they may not be optimal for all membrane proteins and protein–ligand complexes. In addition, when predicting protein–protein complexes in membrane environments, the ranking score should incorporate a protein–protein interface term (e.g., “ipTM_protein–protein_”). Future work will evaluate the optimal parameters using a more diverse set of membrane-protein systems.

### Assessing CoMPLip for ligand-binding pose prediction

To test the generality of CoMPLip for predicting ligand-binding poses in membrane proteins, a benchmark set was prepared by selecting experimentally determined structures of membrane protein–ligand complexes that were not included in the training data of AF3. The dataset comprises 65 structures. The criteria used to construct the dataset are described in the Methods. For each target, structural predictions were performed under two conditions: standard AF3 prediction without lipids and CoMPLip prediction with lipids, in which 100 molecules of 1-monoolein were added. Five seeds were used for each condition. For evaluation, structures with the highest *S*_AF3_ scores were selected for lipid-free predictions, whereas those with the highest *S*_CoMPLip_ scores were selected for CoMPLip predictions. The predicted structures were aligned to the corresponding experimental structure using the protein Cα atoms, and the ligand RMSDs relative to the experimental ligand poses were calculated for all targets. The ligand-binding pose was considered correct when the ligand RMSD was < 4 Å.

**Table 1.** Comparison of ligand-pose prediction accuracy between standard AF3 and CoMPLip. Results for a benchmark dataset of 65 membrane protein–ligand complexes. Predicted ligand-binding poses were classified as correct (ligand RMSD < 4 Å) or incorrect (≥ 4 Å) relative to the experimental ligand poses.

|  | Count |
| --- | --- |
| Correct (w/o) | 41 |
| Improved by CoMPLip | 4 |
| Incorrect (w/o, w/) | 20 |
| Total | 65 |

Correct ligand poses were predicted without lipids for 41 of the 65 targets. Among the remaining 24 targets, CoMPLip improved the predictions for 4 targets, yielding correct poses only when lipids were included (Fig. 4A–D). For the remaining 20 targets, incorrect poses were predicted under both conditions. Conversely, among the 41 targets for which correct poses were predicted without lipids, the CoMPLip predictions produced incorrect poses for five targets (Supplementary Fig. 3A–E). In one case, CoMPLip predicted a different conformation of the membrane protein from that observed in the experimental structure (Supplementary Fig. 3E).

**Figure 4.**
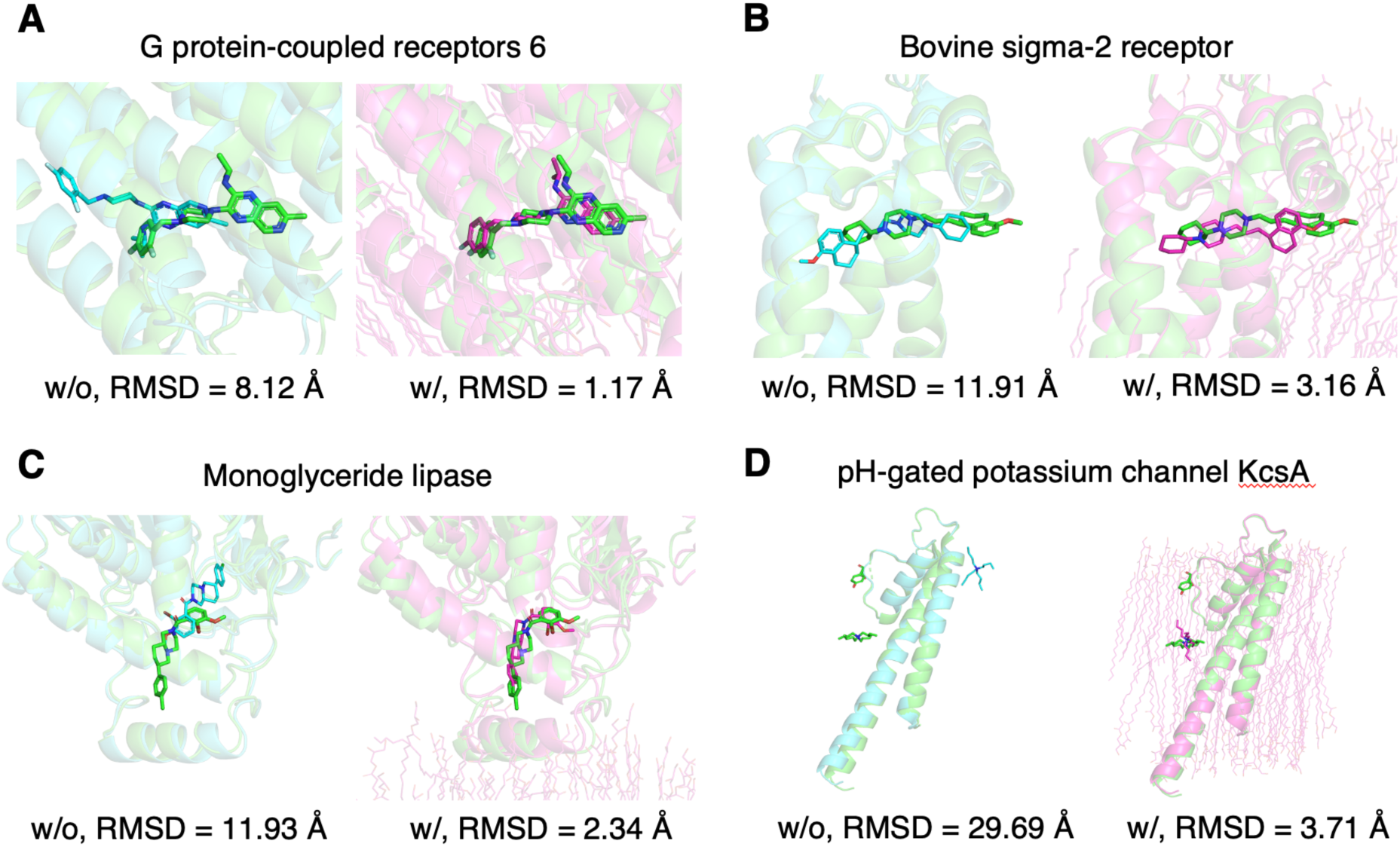
Examples of improved ligand-binding pose predictions with CoMPLip. The experimental structures are shown in green, the predicted structures without lipids (w/o) are shown in cyan, and the CoMPLip predictions with 100 molecules (w/) are shown in magenta. The ligand RMSD values for each prediction are indicated below. **A** G protein-coupled receptor 6 (PDB ID: 8T1V). **B** Bovine sigma-2 receptor (PDB ID: 7M93). **C** Monoacylglycerol lipase (PDB ID: 7ZPG). **D** Potassium channel, KcsA (PDB ID: 8THN).

### Improved domain separation in single-pass membrane proteins

Next, we addressed challenge (ii), the membrane-imposed separation of extracellular domains (ECDs) and intracellular domains (ICDs) across the transmembrane (TM) region in full-length single-pass membrane proteins. The full-length structures of 123 single-pass proteins (Table S2) were predicted using the standard AF3 prediction (without lipids) and CoMPLip prediction (with 50 molecules of 1-monoolein), with five seeds used for each condition. For these proteins, the experimentally determined structures of the ECD, TM region, and ICD were available individually and had been solved before the AF3 training cutoff. However, to the best of our knowledge, no experimentally determined full-length structures encompassing all the three domains are available. Therefore, these targets provide a test of the ability of AF3 to assemble a correct full-length ECD–TM–ICD architecture without relying on full-length structural templates available before the training cutoff. For each target protein, the structures with the highest *S*_AF3_ and *S*_CoMPLip_ scores were selected for lipid-free and CoMPLip predictions, respectively, and domain separation was assessed visually.

A clear separation between ECD and ICD across the TM region was observed in 20 of the 123 models predicted without lipids (Table 2). In contrast, clear domain separation was observed in 61 of the 123 models predicted with CoMPLip (representative structures are shown in Fig. 5). In the CoMPLip prediction, lipid molecules were consistently placed around the TM helix, forming a bilayer-like lipid environment. This lipid layer surrounding the TM helix appears to prevent direct contact between ECD and ICD. However, in some of the predicted structures, the ECD and ICD approached each other by bypassing the lipid layer (Supplementary Fig. 4). This behavior may arise when a limited number of lipid molecules form an incomplete bilayer-like structure with gaps around the TM helix. Increasing the number of lipid molecules may reduce such gaps, thereby promoting ECD–ICD separation.

**Figure 5.**
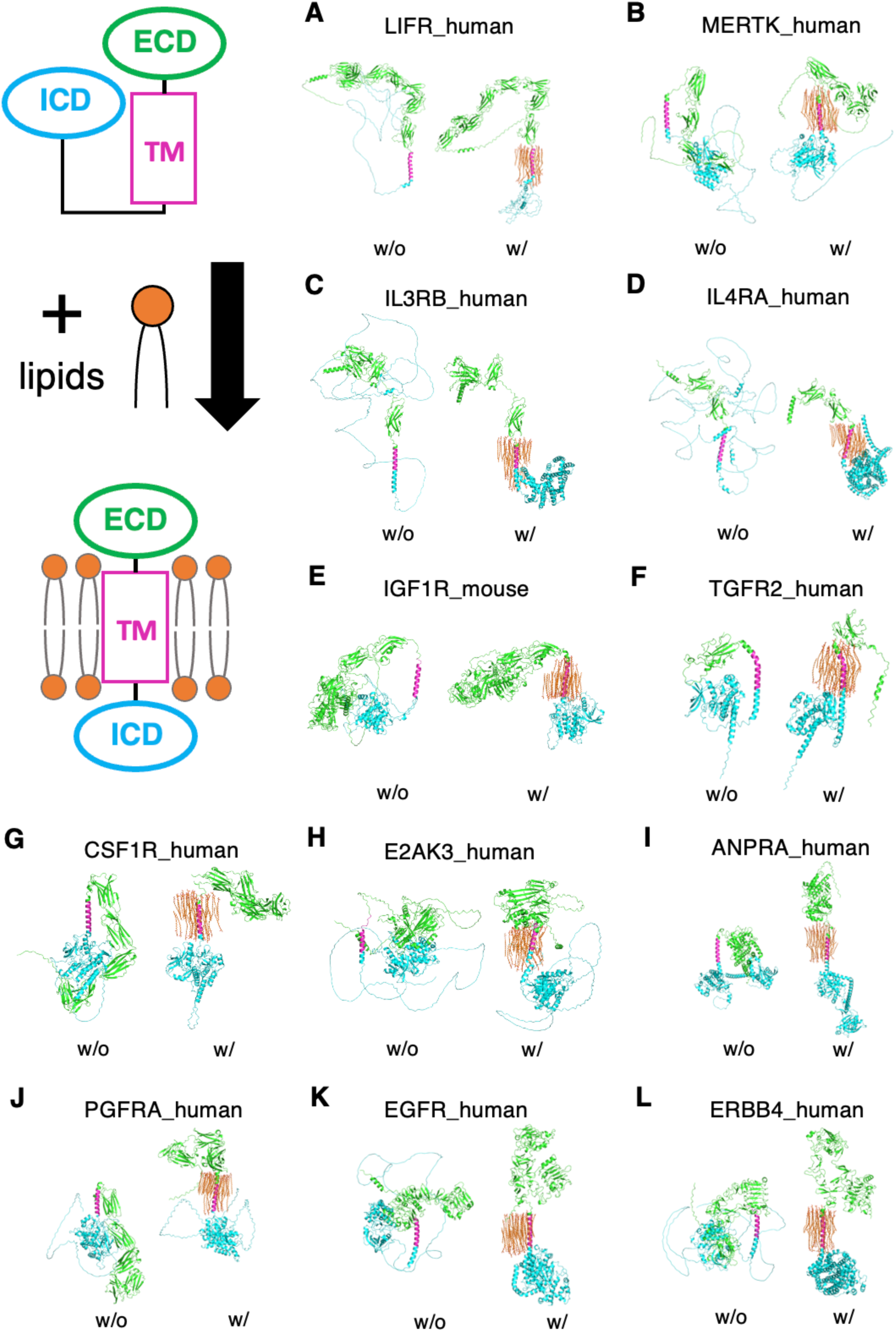
Examples of full-length single-pass transmembrane protein predictions by standard AF3 and CoMPLip. **A–L**: Representative predictions for full-length single-pass transmembrane proteins. The extracellular domain (ECD) is shown in green, the intracellular domain (ICD) in cyan, and the transmembrane (TM) region in magenta. Added lipids are shown as orange sticks. Predicted structures without lipids (standard AF3) are shown on the left and CoMPLip predictions with 50 molecules of 1-monoolein are shown on the right.

**Table 2.**
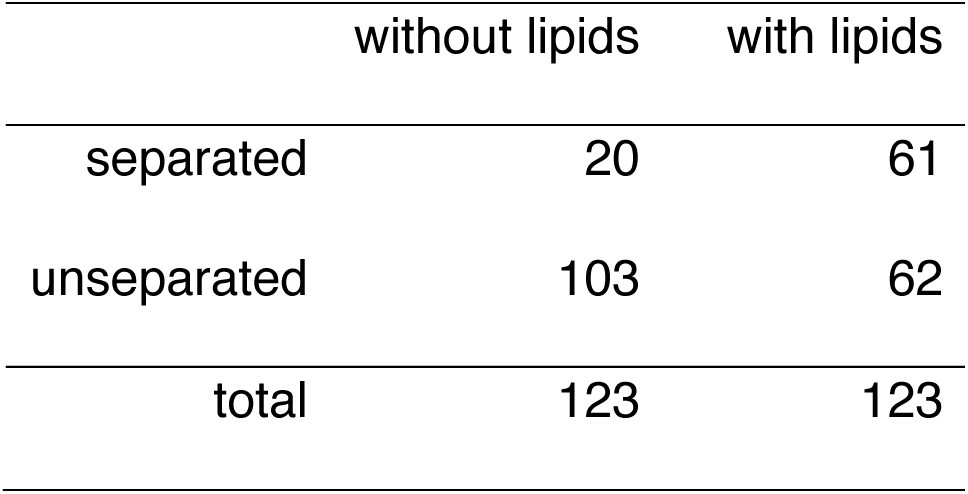
Comparison of domain separation predictions between standard AF3 and CoMPLip. Contingency table for 123 single-pass transmembrane proteins classified as separated or unseparated between the extracellular and intracellular domains across the transmembrane region.

|  | without lipids | with lipids |
| --- | --- | --- |
| separated | 20 | 61 |
| unseparated | 103 | 62 |
| total | 123 | 123 |

To further examine the effect of CoMPLip, full-length structures of the human epidermal growth factor receptor (EGFR) were predicted for both monomer and dimer forms (Fig. 6). For dimer prediction, two epidermal growth factor molecules were included to mimic EGFR activation. For both monomers and dimers, domain separation was improved by CoMPLip using POPC lipids that resembled the native membrane environment. Interestingly, the conformational states of the ECD and ICD (kinase domains) in the predicted structures were mutually incompatible ^18^. In the CoMPLip prediction of the EGFR monomer prediction, ECD consistently adopted an inactive conformation, whereas ICD adopted an active conformation (Fig. 6). Similarly, in the predicted EGFR dimer, the ECDs adopted an active conformation, whereas the ICDs formed an inactive-like symmetric dimer (Fig. 6 and Supplementary Fig. 5). When incompatible domain conformations are obtained, one possible strategy is to guide the prediction using experimentally determined active or inactive structures as templates, thereby biasing the ECD and ICD toward compatible conformational states.

**Figure 6.**
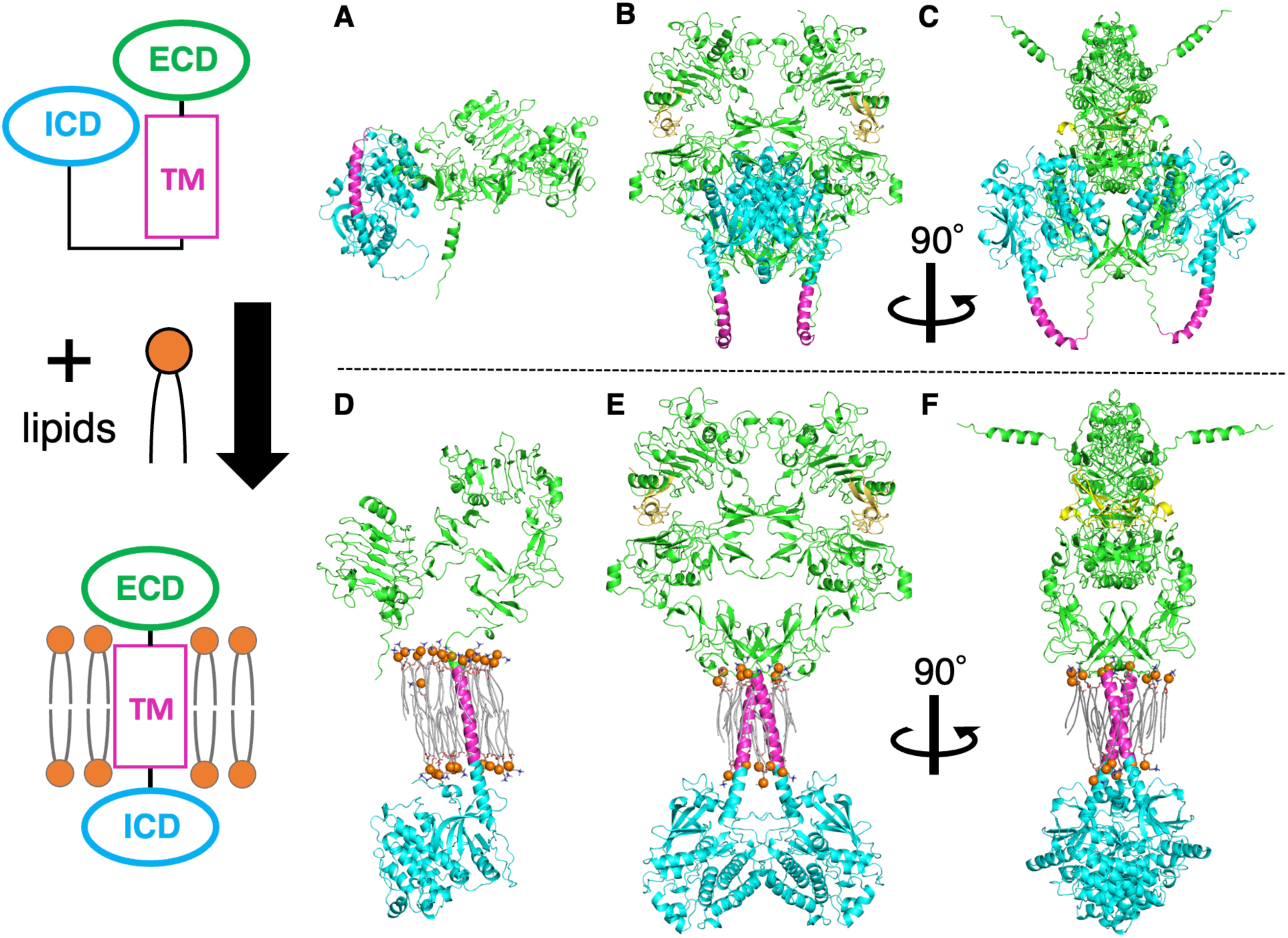
Full-length structures of human EGFR monomer and EGFR–EGF dimer predicted by standard AF3 (without lipids) and by CoMPLip with POPC lipids. The extracellular domain (ECD) is shown in green, intracellular kinase domain (ICD) in cyan, transmembrane helix in magenta, EGF in yellow, POPC headgroups in orange, and POPC acyl chains in gray. **A** Monomeric EGFR predicted without lipids. **B** EGFR–EGF dimer predicted without lipids. **C** Side view of panel B. **D** Monomeric EGFR predicted with POPC lipids. **E** EGFR–EGF dimer predicted with POPC lipids. **F** Side view of panel E.

### Improved conformational sampling with CoMPLip

Finally, we addressed challenge (iii), the conformational sampling of dynamic membrane proteins, by assessing the effect of CoMPLip on the diversity of the predicted conformations. As a test case, we used the sodium taurocholate co-transporting polypeptide (NTCP), a transmembrane transporter that undergoes large conformational changes. The outward-facing (OF) and inward-facing (IF) structures of human NTCP have both been determined experimentally ^19,20^, but neither conformation has been included in the AF3 training set ^7^. NTCP predictions were performed under two conditions: without lipids and with lipids (CoMPLip with 100 1-monoolein molecules). Predicted models were classified as OF or IF based on the RMSD of Cα atoms relative to the experimental structures (PDB ID: 7FCI for OF and 7PQG for IF), using a threshold of RMSD < 2 Å. In the standard AF3 prediction without lipids, all 500 structures adopted the IF conformation (RMSD < 2 Å to 7PQG). In contrast, in CoMPLip predictions, 305 models adopted the IF conformation and 195 models adopted the OF conformation (Fig. 7). This difference is also evident in the position of TM8b (Fig. 8), which contributed to the opening and closing of the central pore. To further investigate the structural diversity of the CoMPLip models, the distance between Cα atoms of L31 in TM1 and Q264 in TM8b was calculated (Supplementary Fig. 6). The predicted structures were divided into two clusters, corresponding to the IF and OF conformations. In the CoMPLip model, which was closest to the OF experimental structure, a lipid molecule was positioned near TM8b, which would sterically clash with the TM8b position in the IF structure (Supplementary Figs. 7A and 7B). In contrast, some OF-like models lacked lipids in the immediate vicinity of TM8b, where steric overlap would otherwise be expected (Supplementary Fig. 7C and 7D). These observations suggest that lipid placement near TM8b varies across models and does not show a consistent relationship with the IF/OF classification. Thus, although CoMPLip can generate both IF- and OF-like structures, no clear correlation was observed between the predicted lipid positions and the emergence of a particular conformational state. These results suggest that introducing an explicit membrane environment using CoMPLip can enhance the conformational sampling of dynamic membrane transporters.

**Figure 7.**
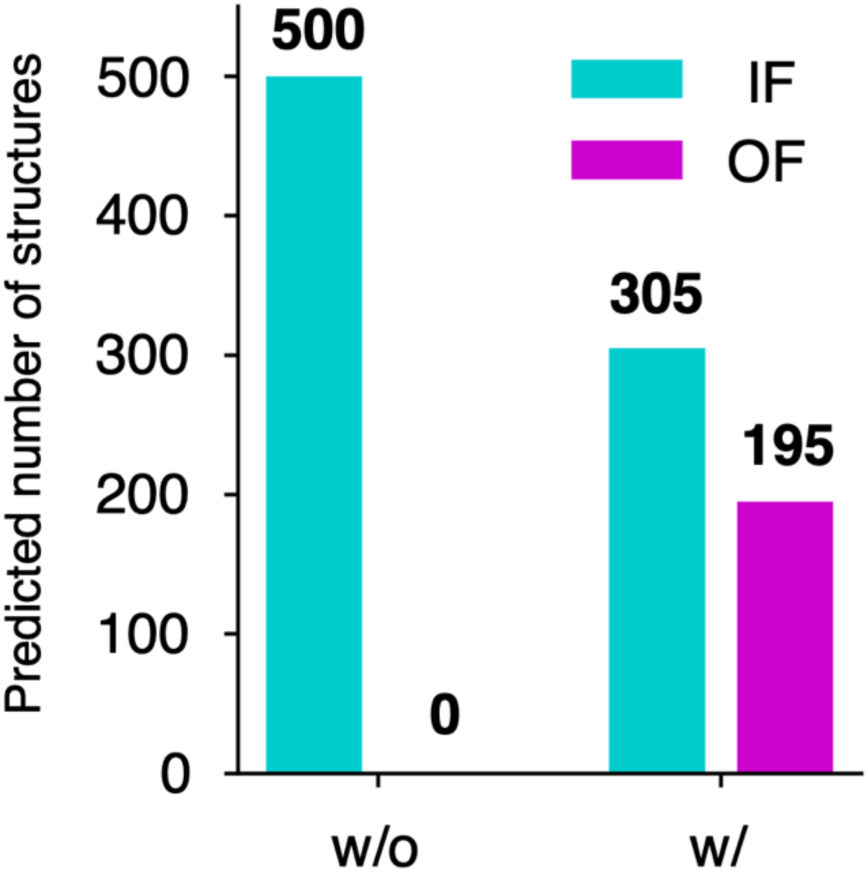
Enhanced conformational sampling of NTCP using CoMPLip. Predictions for NTCP were performed under two conditions: without lipids (standard AF3) and with 100 1-monoolein (CoMPLip), using 100 seeds (500 predicted structures) per condition. The number of inward-facing (IF) and outward-facing (OF) conformations are represented in cyan and magenta, respectively.

**Figure 8.**
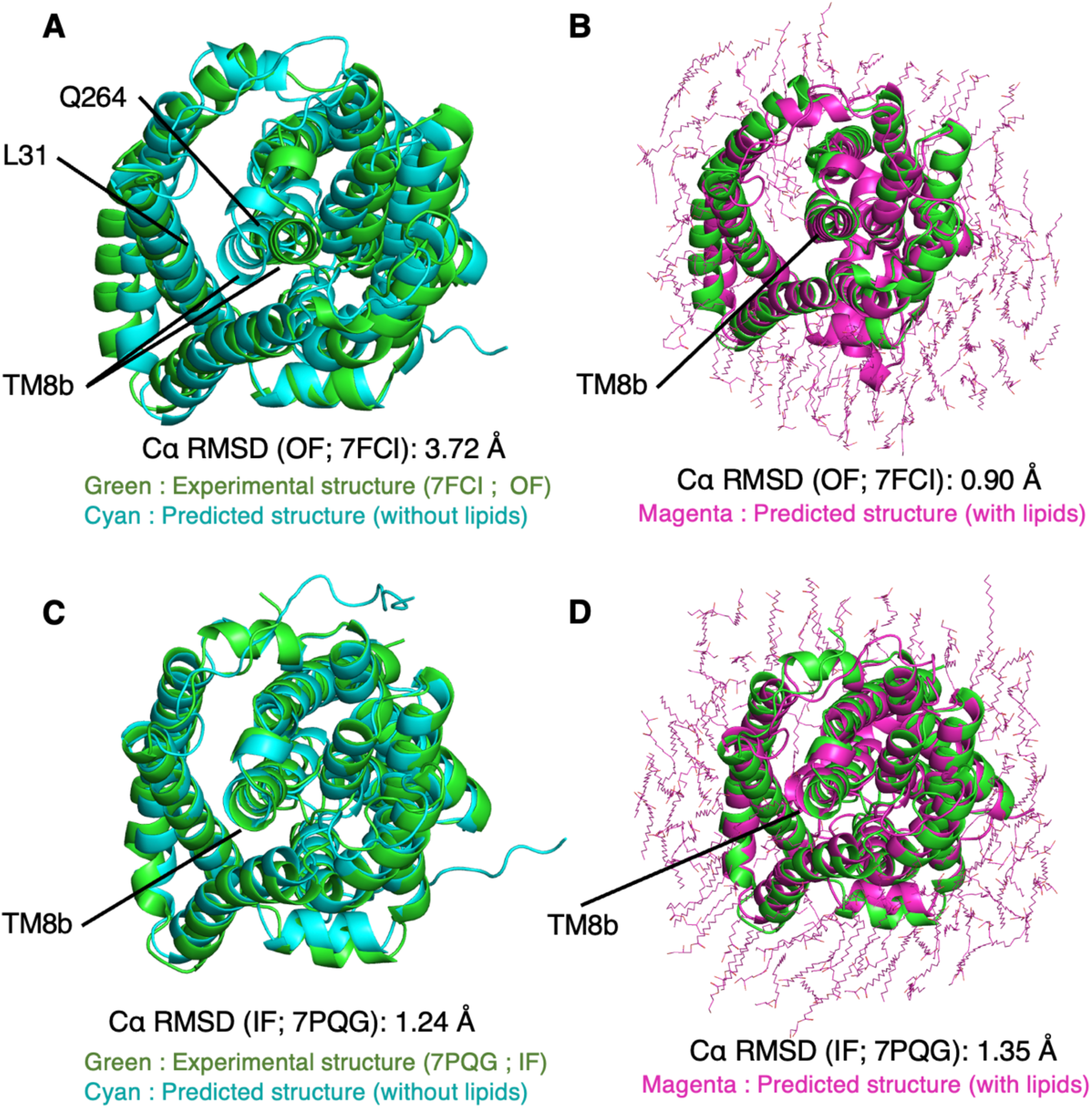
Lipid addition enables sampling of both inward- and outward-facing NTCP conformations. **A** Without lipids (100 seeds), the predicted structure (cyan) is closest to the outward-facing reference structure (green; PDB ID: 7FCI). The RMSD of Cα atoms is 3.72 Å. **B** With lipids (CoMPLip, 100 seeds), the predicted structure (magenta) closest to the outward-facing reference is shown (RMSD = 0.90 Å). **C** Without lipids, the predicted structure (cyan) closest to the inward-facing reference (green; PDB ID: 7PQG) is shown (RMSD = 1.24 Å). **D** With lipids, the predicted structure (magenta) closest to the inward-facing reference is shown (RMSD = 1.35 Å).

### Summary, limitations, and outlook

In summary, we introduced CoMPLip, a simple approach for improving AlphaFold 3 structure prediction for membrane proteins by co-folding lipid molecules. Using CoMPLip, we demonstrated improvements in the three challenges outlined in the Introduction: (i) improved ligand-binding poses in membrane proteins, (ii) improved separation between the extracellular and intracellular domains in full-length single-pass membrane proteins, and (iii) enhanced sampling of multiple conformations. We further developed *S*_CoMPLip_, a CoMPLip-specific ranking score that focuses on the prediction accuracy for the target protein and ligand rather than the added lipids.

The coefficients of *S*_CoMPLip_ were optimized using the RseP–BAT system. Additionally, the lipid copy number and carbon chain length were optimized. As a practical guideline, CoMPLip requires a sufficient number of lipid molecules to cover the transmembrane region during prediction. For example, co-folding approximately 100 molecules of 1-monoolein may provide a reasonable starting point for a four-pass membrane protein.

A limitation of CoMPLip is that co-folding lipid molecules requires more GPU memory than the standard AF3 predictions, which can make structure prediction of large membrane proteins with sufficient lipids difficult. In the benchmark dataset of 65 membrane protein-ligand complexes, CoMPLip improved the ligand-binding poses for four targets, although the overall success rate of ligand-pose prediction did not increase. These results suggest that further optimization of the ranking score and CoMPLip parameters is necessary. In NTCP predictions, lipid insertion into the pore region was also observed. Notably, the distribution of the predicted lipid molecules cannot be directly controlled and is instead determined by the model, which can lead to undesired lipid positioning (e.g., within the pore). Including substrates may help reduce this behavior.

In addition to lipid co-folding in CoMPLip, other small molecules may serve as effective co-input additives and improve the prediction accuracy using the same protocol. Taken together, CoMPLip, combined with appropriate lipid additives and an adapted ranking score, provides an easily implemented protocol for improving membrane protein structure prediction and may facilitate structure-based studies of challenging transmembrane targets. By introducing an explicit membrane context into structure prediction, CoMPLip provides a practical framework for predicting membrane-protein structures in biologically relevant environments.

## Methods

### AlphaFold 3 settings and computational environment

In the CoMPLip structure predictions, AF3 was run using SMILES strings for lipid molecules, in addition to the input information for the target membrane proteins and ligands. All structural predictions in this study were performed using AF3, version 3.0.1, with the JSON input schema version 2. For each target, the number of random seeds was set to either 5 or 100, as indicated, and the number of diffusion samples per seed was set to five. Structural predictions for all targets, except the full-length EGFR structures, were performed using NVIDIA A100 GPUs (40 GB). Because the CoMPLip-based EGFR dimer prediction required substantially larger GPU memory, the system was predicted using an NVIDIA H100 GPU (94 GB). The number of lipids included in each CoMPLip prediction was selected within the token limit allowed by the GPU used. Unless otherwise noted, AF3 was run using the default model parameters provided in version 3.0.1. All structures were visualized and inspected using PyMOL (Schrödinger LLC, USA).

### Structure predictions of complex of RseP and BAT

The amino acid sequence of *Escherichia coli* RseP was obtained from PDB entry 7W6X ^15^. The target ligand, batimastat (BAT), was specified by the following SMILES string obtained from a PDB ^21^ entry:

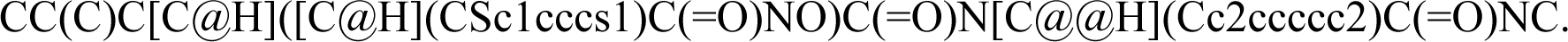

The Zn²⁺ ion in the active site was specified by the PDB chemical component dictionary (CCD) code “ZN.” In the CoMPLip predictions for RseP, 1-monoolein (glyceryl monooleate) was used as the added lipid and was specified by the SMILES obtained from PubChem ^22^:

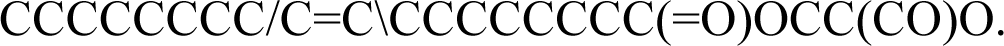

In the JSON input files, each backslash (\) in the SMILES string was escaped as \\. An example of the AF3 JSON input used for CoMPLip is shown in Supplementary Fig. 8.

For lipid copy-number analysis, the number of 1-monoolein molecules was increased from 0 to 100 in increments of 10 in the CoMPLip-based RseP–BAT complex prediction, and the ligand RMSD was measured for all predicted structures. For ligand RMSD calculations, predicted structures were aligned to the experimental RseP structure (PDB ID: 7W6X) using Cα atoms of the transmembrane segments (residues 2–33, 94–122, 376–415, 423–447) ^15^. After alignment, the BAT poses in the predicted and experimental structures were compared and the RMSD of the non-hydrogen atoms was calculated using pyDockRMSD ^23^. For the carbon chain length analysis, we used a series of lipids sharing the glycerol backbone of 1-monoolein with different acyl-chain lengths (general structure: R–C(=O)O–CH₂–CH(OH)–CH₂OH; *n* = 1–17, where *n* denotes the number of carbons in the alkyl chain R and R = CH₃(CH₂)*_n_*_−1_). For each carbon chain length (*n* = 1–17), CoMPLip prediction was performed using 100 lipid molecules. The prediction without lipids was also included as *n* = 0 for comparison.

### Structure predictions of other proteins and ligands

To prepare the ligand-binding transmembrane protein test set, transmembrane protein–ligand complexes were collected from the RCSB PDB ^21^ according to the following criteria: (i) released from October 2021, and therefore not included in the AF3 training data ^7^; (ii) containing at least one ligand with molecular weight ≥ 100; (iii) determined at a resolution ≤ 3.0 Å; (iv) containing ≤ 800 amino acid residues; and (v) annotated in OPM ^24^ or PDBTM ^25^. For the collected PDB entries, all non-protein molecules, including lipids, detergents, and crystallization additives, were removed except for the target ligand. The final set consisted of 65 targets for which AF3 predictions were successfully completed (Supplementary Table 1 and Supplementary Data 1). For each target, predicted structures were aligned to the corresponding experimental structure using protein Cα atoms, ligand RMSD of non-hydrogen atoms was computed using pyDockRMSD ^23^.

### Structure predictions of single-pass transmembrane proteins

To prepare single-pass transmembrane protein test set, UniProt ^26^ was queried for proteins satisfying the following criteria: (i) annotated with the subcellular location “single-pass membrane protein”; (ii) containing ≤ 1400 amino acid residues; (iii) having more than 100 residues on both side of the TM segment; (iv) reviewed (UniProtKB/Swiss-Prot); (v) having at least one recently released experimental structure with a PDB ID starting with 8 or 9. The final set consisted of 123 proteins for which AF3 predictions were successfully completed (Supplementary Table S2 and Supplementary Data 2). For each target, structural prediction was performed under two conditions, without lipids and with 50 molecules of 1-monoolein, using five seeds for each condition.

For EGFR structure prediction, the human EGFR sequence (UniProt ID: P00533) corresponding to residues 1–1022 was used. For CoMPLip predictions, 15 or 30 POPC molecules were added. The SMILES of the POPC were obtained from the RCSB PDB:

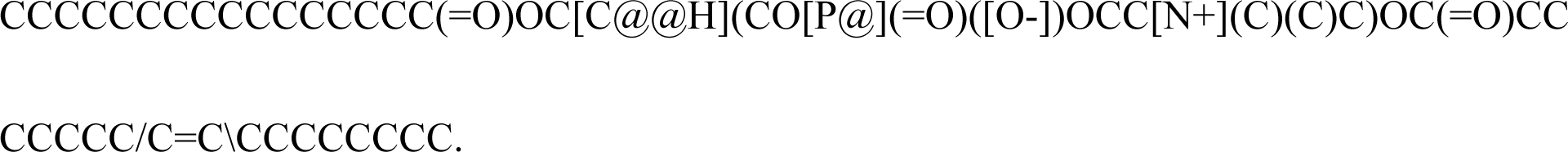

### Structure predictions of NTCP

For CoMPLip prediction of human NTCP, the UniProt sequence Q14973 was used and 100 molecules of 1-monoolein were added. For comparison, the prediction without lipids was also performed. Both predictions were run with 100 seeds. For all 500 predicted models in each condition, RMSDs of Cα atoms relative to the inward-facing (IF) structure (PDB ID: 7PQG) and outward-facing (OF) structure (PDB ID: 7FCI) were calculated. Predicted structures with RMSD < 2 Å to either reference structure was classified as belonging to that conformation.

## Acknowledgements

This research was supported by the Platform Project for Supporting Drug Discovery and Life Science Research (Basis for Supporting Innovative Drug Discovery and Life Science Research (BINDS)) from the Agency for Medical Research and Development (AMED) under grant number JP25ama121023 (Mi. I.), and by AMED under grant number JP25fk0310525 (Mi. I.). This work was supported by JST SPRING, Japan Grant Number JPMJSP2179 (H. O.). This work used computational resources of the TSUBAME4.0 provided by Tokyo Institute of Technology through BINDS from AMED (JP25ama121023 (Mi. I.)). This study used computational resources from the Yokohama City University, Tsurumi Campus, Japan.

## Author contributions

H. O. conceived and designed the study, performed the computations, curated the data, analyzed the results, prepared the figures, and wrote the manuscript. Ma. I., T. E., T. Y., and Mi. I. reviewed and edited the manuscript. Mi. I. supervised the study.

## Competing Interests

The authors declare no competing interests.

## Notes

### Competing Interest Statement

The authors have declared no competing interest.

