## Supplementary Information for "Co-folding of Membrane Proteins and Lipid Molecules Improves Membrane-Protein Structure Prediction Accuracy"

**Contents**
**Supplementary Fig. 1 – 8**
**Supplementary Table 1, 2**
**Supplementary Data 1, 2**
**Supplementary References**

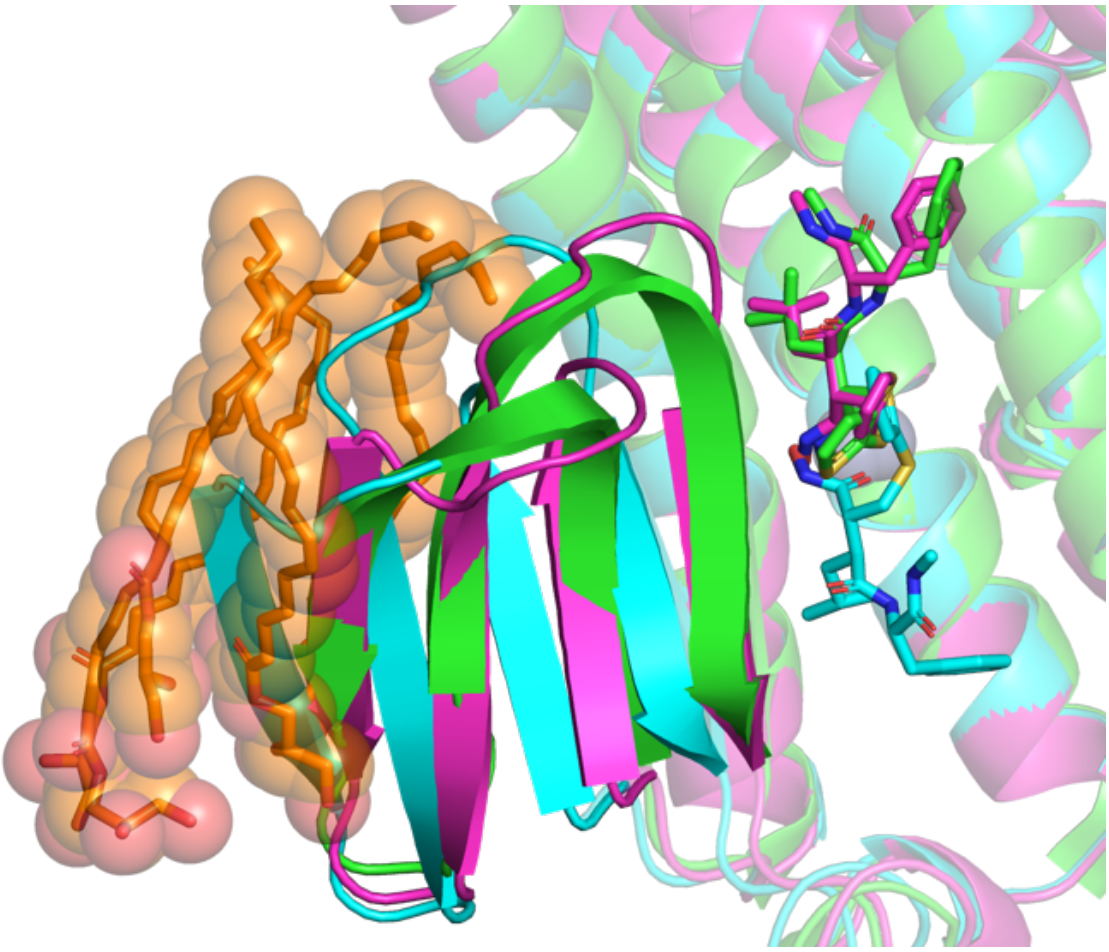

**Supplementary Fig. 1 | Predicted MREβ region and associated lipid molecules.** Green, experimental structure of RseP (PDB ID: 7W6X); cyan, standard AF3 prediction (w/o lipids); magenta, CoMPLip prediction (w/ lipids); orange, 1-monoolein molecules in the CoMPLip prediction located near the MREβ region. Structures were aligned using Cα atoms of the transmembrane region of the experimental structure. Lipid molecules predicted near the MREβ region are highlighted as spheres.

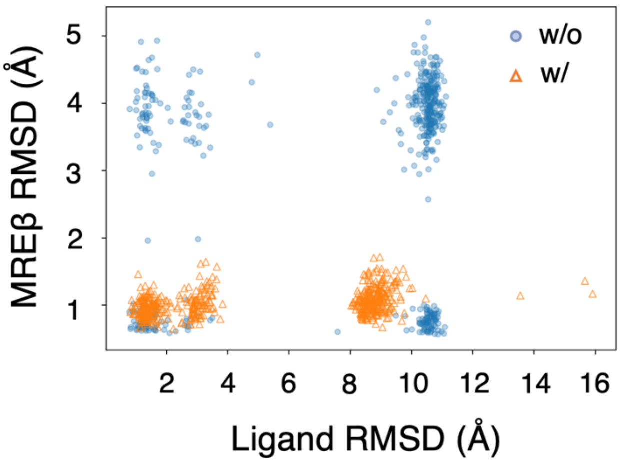


**Supplementary Fig. 2 | RMSD distributions of the MREβ region and ligand across predicted structures of RseP–BAT complex.**

Predicted structures were plotted as ligand (BAT) RMSD versus MREβ RMSD. Standard AF3 predictions (w/o lipids) are represented as blue circles, and CoMPLip predictions (w/ lipids) as orange triangles (n = 500 each).

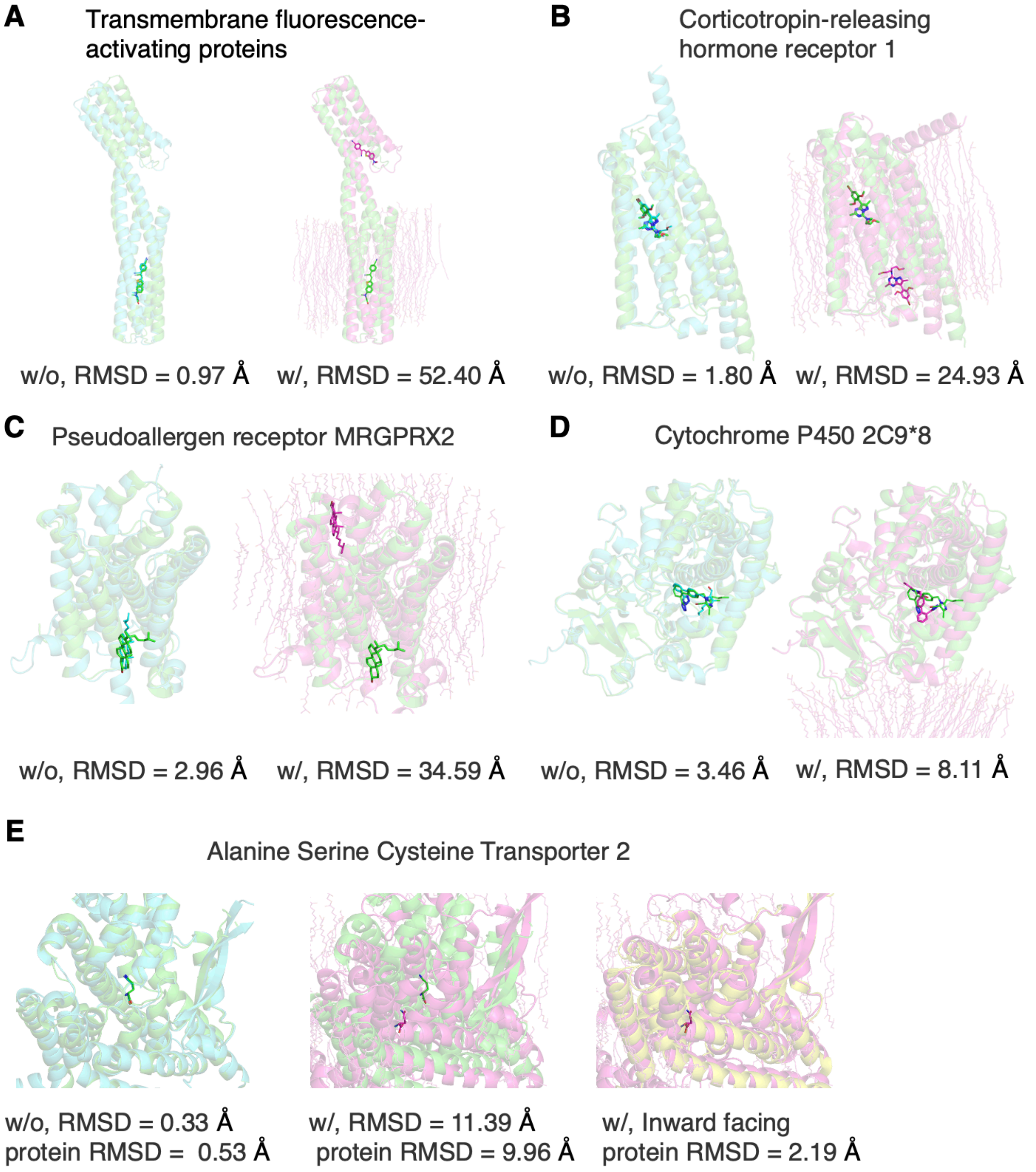


**Supplementary Fig. 3 | CoMPLip-induced changes in ligand prediction.**
Experimental structures are shown in green, standard AF3 predictions (without lipids) in cyan, and CoMPLip (with 100 lipids) in magenta. The ligand RMSD values were as follows: **A** Transmembrane fluorescence-activating proteins (PDB ID: 8T1V). **B** Corticotropin-releasing hormone receptor 1 (PDB ID: 8GTG). **C** Pseudoallergen receptor MRGPRX2 (PDB ID: 7VV4). **D** Cytochrome P450 2C9*8 (PDB ID: 7RL2). **E** Alanine Serine Cysteine Transporter 2 (PDB ID: 8QRP, green, outward-facing; PDB ID: 6RVX, yellow, inward-facing).

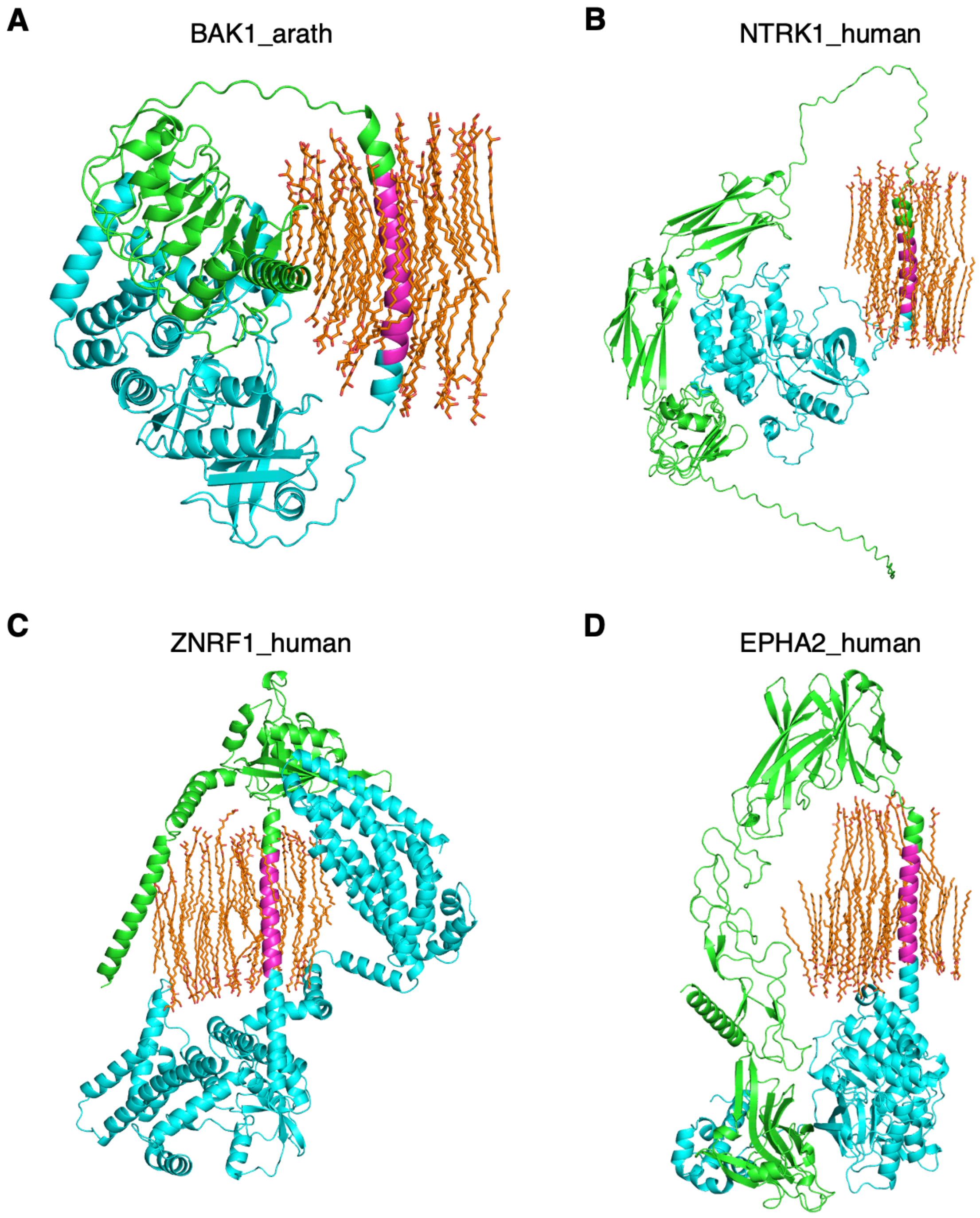


**Supplementary Fig. 4 I Examples where extracellular and intracellular domains are predicted to contact across the transmembrane region.**

**A–D**: N-terminal domains are shown in green, C-terminal domains in cyan, and transmembrane domains in magenta. The lipids are shown as orange sticks.

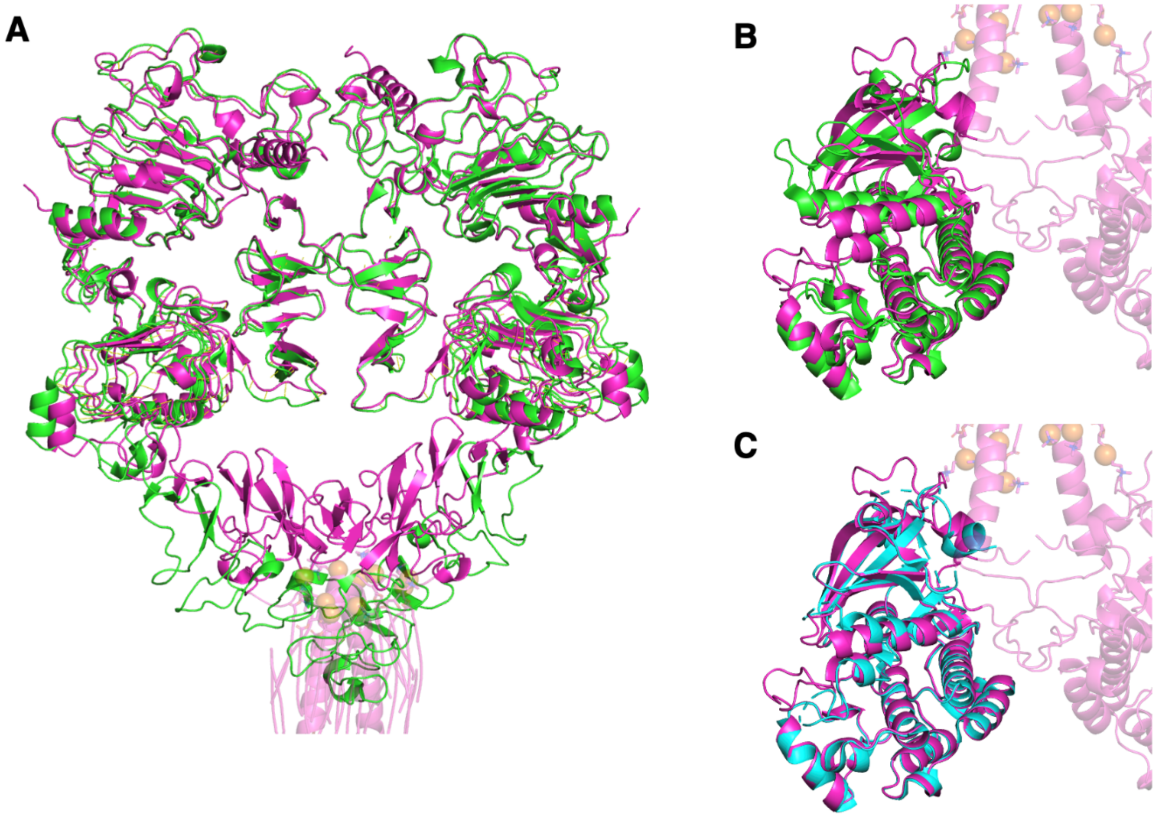


**Supplementary Fig. 5 | Comparison of the predicted EGFR–EGF dimer with experimental structures.**

**A** Alignment of the extracellular domain (ECD) of the predicted EGFR–EGF dimer (magenta) with the ECD of active EGFR (PDB ID: 7SYD; green). RMSD = 1.614 Å. **B** Alignment of the intracellular kinase domain (ICD) of the predicted EGFR–EGF dimer (magenta) with the ICD of the active EGFR (PDB ID: 1M14; green). RMSD = 1.636 Å. **C** Alignment of the predicted ICD (magenta) with the ICD of the inactive EGFR (PDB ID: 1XKK; cyan). RMSD = 0.391 Å.

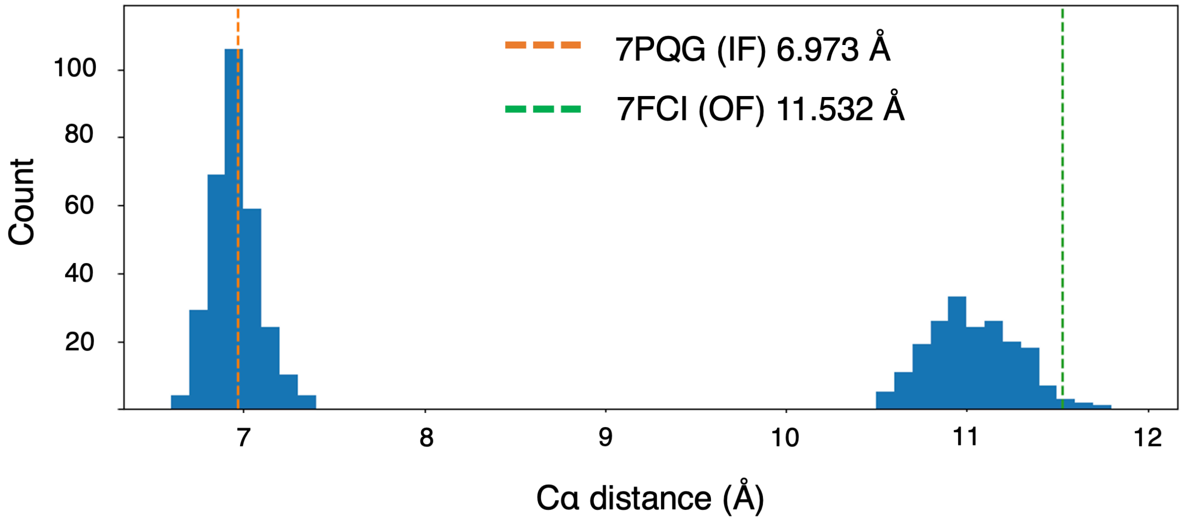


**Supplementary Fig. 6 | Distribution of the L31–Q264 Cα–Cα distance in CoMPLip-predicted NTCP structures.**

Distances were calculated for 500 CoMPLip-predicted structures and plotted as histograms (bin width, 0.1 Å). The orange and green dashed lines indicate the inward-facing (PDB ID: 7PQG) and outward-facing (PDB ID: 7FCI) experimental structures, respectively.

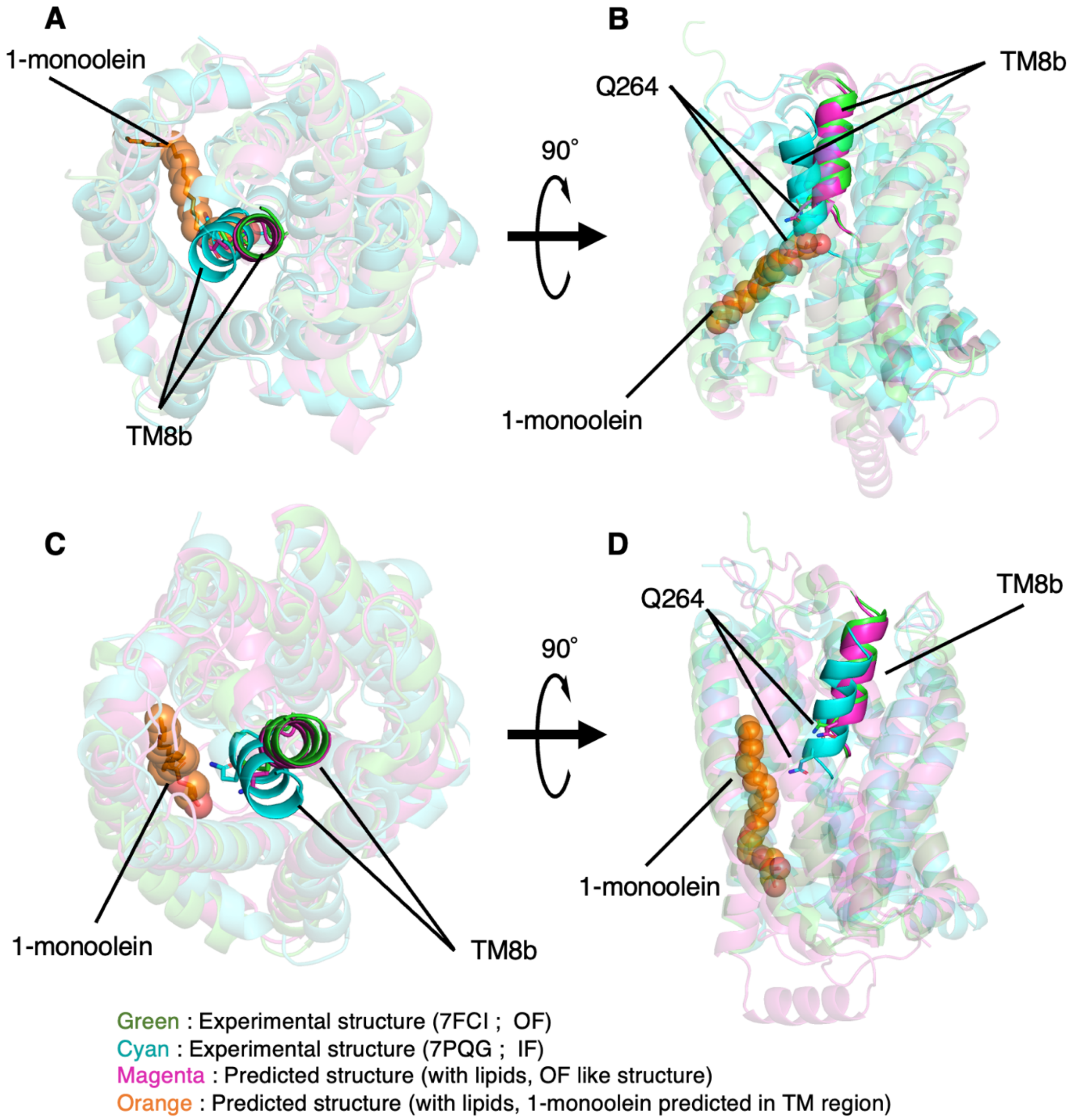


**Supplementary Fig. 7 l Lipid placement near TM8b in CoMPLip predictions.** Predicted NTCP structures (magenta) were aligned with the outward-facing (OF) and inward-facing (IF) experimental structures (PDB IDs: 7FCI (OF, green) and 7PQG (IF, cyan)) using Cα atoms in the transmembrane region. Lipid molecules closest to TM8b are shown as orange spheres. **A** Top view of the prediction closest to the OF structure. **B** Side view of (A). **C** Example of a prediction in which lipids are relatively distant from TM8b. **D** Side view of (C).

```
{
    "name": "RseP_100crystal_Zn_Bat",
    "modelSeeds": [1,2,3,4,5],
    "sequences": [
        {
            "protein": {
                "id": "A",
                "sequence": "MLSFLWDLASFIVALGVLITVHEFGHFWVARRCGVRVERFSIGFGKALWRRTDKLGTEYVIALIPLGGYVKML
DERAEPVVPELRHHAFNNKSVGQRAAIIAAGPVANFIFAIFAYWLVFIIGVPGVRPVVGEIAANSIAAEAQIAPGTELKAVDGIETPDWDAVRLQLVDKIGD
ESTTITVAPFGSDQRRDVKLDLRHWAFEPDKEDPVSSLGIRPRGPQIEPVLENVQPNSAASKAGLQAGDRIVKVDGQPLTQWVTFVMLVRDNPGKSLALEIE
RQGSPLSLTLIPESKPGNGKAIGFVGIEPKVIPLPDEYKVVRQYGPFNAIVEATDKTWQLMKLTVSMLGKLITGDVKLNNLSGPISIAKGAGMTAELGVVYY
LPFLALISVNLGIINLFPLPVLDGGHLLFLAIEKIKGGPVSERVQDFCYRIGSILLVLLMGLALFNDFSRLGTENLYFQ"
            }
        },
        {
            "ligand": {
                "id": ["WE", "WF", "WG", "WH", "WI", "WJ", "WK", "WL", "WM", "WN", "WO", "WP", "WQ", "
WR", "WS", "WT", "WU", "WV", "WW", "WX", "WY", "WZ", "XA", "XB", "XC", "XD", "XE", "XF", "XG", "XH", "
XI", "XJ", "XK", "XL", "XM", "XN", "XO", "XP", "XQ", "XR", "XS", "XT", "XU", "XV", "XW", "XX", "XY", "
XZ", "YA", "YB", "YC", "YD", "YE", "YF", "YG", "YH", "YI", "YJ", "YK", "YL", "YM", "YN", "YO", "YP", "
YQ", "YR", "YS", "YT", "YU", "YV", "YW", "YX", "YY", "YZ", "ZA", "ZB", "ZC", "ZD", "ZE", "ZF", "ZG", "
ZH", "ZI", "ZJ", "ZK", "ZL", "ZM", "ZN", "ZO", "ZP", "ZQ", "ZR", "ZS", "ZT", "ZU", "ZV", "ZW", "ZX", "
ZY", "ZZ"],
                "smiles": "CCCCCCCC/C=C\\CCCCCCCC(=O)OCC(CO)O"
            }
        },
        {
            "ligand": {
                "id": "B",
                "smiles": "CC(C)C[C@H]([C@H](CSc1cccs1)C(=O)NO)C(=O)N[C@@H](Cc2ccccc2)C(=O)NC"
            }
        },
        {
            "ligand": {
                "id": "C",
                "ccdCodes": ["ZN"]
            }
        }
    ],
    "dialect": "alphafold3",
    "version": 2
}

(END)
```


**Supplementary Fig. 8 | Example of an AlphaFold 3 input file.**

Representative JSON-formatted input used for AF3 structure prediction is shown.

**Supplementary Table 1 | PDB IDs of 65 ligand-bound membrane proteins[1].**
6zfz, 6zg4, 7f61, 7f8u, 7khm, 7m93, 7mdc, 7pp1, 7q0m, 7r1k, 7rl2, 7rox, 7spt, 7srq, 7ss6, 7sus, 7ufb, 7ug0, 7ul2, 7um4, 7voe, 7vv4, 7w41, 7wc6, 7xrr, 7zg9, 7zpg, 8c9w, 8cic, 8cu6, 8de4, 8do0, 8du3, 8e7w, 8ex4, 8fyn, 8gne, 8gtg, 8j1n, 8jsw, 8jt8, 8jt9, 8jtb, 8ovv, 8oyg, 8pwn, 8pz4, 8q2o, 8qct, 8qrp, 8qt6, 8rw0, 8t1v, 8t69, 8thn, 8ubw, 8uf6, 8urp, 8w4b, 8y2d, 9as7, 9d68, 9fup, 9h76, 9ivk

**Supplementary Table 2 | UniProt IDs of 123 single-pass membrane proteins[2].**
Q94JS0, Q9CR68, P13272, Q7Z6A9, Q86YW5, Q6WZB0, P0A221, O75381, O60667, P15751, Q9NZA1, P08138, O14763, P19438, O43464, P16872, P20333, Q01114, P58335, P36897, P19235, Q04771, O00481, Q8WVV5, Q7KYR7, Q8CGK5, Q4F883, Q9HBE5, Q92692, Q0PMD2, A4YDT1, P37173, P04843, Q94F62, Q6UD73, Q08351, Q5VWK5, P08195, P40238, Q02297, P9WI79, Q60837, O95202, P01833, Q68DV7, O60603, O94901, P04629, P22455, P22607, P21802, Q16620, P11362, P78536, P24394, P22223, Q14162, Q8NFZ4, Q9QUK6, O00206, A0A385DV85, P19550, Q9NZ94, F4KF14, P04624, P04578, O60602, Q9Q714, Q99665, P12489, O15146, P97378, P12830, P32927, P06436, P06437, O15455, Q9P0K1, P13201, Q08345, Q00560, P40189, Q9ULT6, Q01974, Q26261, Q7Z3J2, O70458, P07333, P10721, P29317, O75460, P54764, Q12866, O09127, Q9UIQ6, Q9NR96, Q9NR97, Q8VZG8, P29323, P16066, P25092, P16234, P42702, Q9NZJ5, Q14126, P48356, P48357, Q0JA29, Q9H0H0, P01133, P00533, P04626, P32004, Q6PDJ1, P08575, Q15303, P03726, P08069, P15208, Q60751, P06213, P08581, Q04912

**Supplementary Data**

1. PDB entries and results used for the structure prediction of ligand-bound membrane proteins (Excel, CoMPLip_ligand-binding_protein.xlsx).
2. PDB entries and results used for the structure predictions of single-pass transmembrane proteins (Excel, CoMPLip_single-pass.xlsx).
